# Membrane-mediated forces can stabilize tubular assemblies of I-Bar proteins

**DOI:** 10.1101/2020.06.10.144527

**Authors:** Z. Jarin, A. J. Pak, P. Bassereau, G. A. Voth

## Abstract

Collective action by Inverse-BAR (I-BAR) domains drive micron-scale membrane remodeling. The macroscopic curvature sensing and generation behavior of I-BAR domains is well characterized, and computational models have suggested various mechanisms on simplified membrane systems, but there remain missing connections between the complex environment of the cell and the models proposed thus far. Here, we show a connection between the role of protein curvature and lipid clustering in the stabilization of large membrane deformations. We find lipid clustering provides a directional membrane-mediated interaction between membrane-bound I-BAR domains. Lipid clusters stabilize I-BAR domain aggregates that would not arise through membrane fluctuation-based or curvature-based interactions. Inside of membrane protrusions, lipid cluster-mediated interaction draws long side-by-side aggregates together resulting in more cylindrical protrusions as opposed to bulbous, irregularly shaped protrusions.

**Statement of Significance:** Membrane remodeling occurs throughout the cell and is crucial to proper cellular function. In the cellular environment, I-BAR proteins are responsible for sensing membrane curvature and initiating the formation of protrusions outward from the cell. Additionally, there is a large body of evidence that I-BAR domains are sufficient to reshape the membrane on scales much larger than any single domain. The mechanism by which I-BAR domains can remodel the membrane is uncertain. However, experiments show that membrane composition and most notably negatively-charge lipids like PIP_2_ play a role in the onset of tubulation. Using coarse-grained models, we show that I-BAR domains can cluster negatively charge lipids and clustered PIP_2_-like membrane structures facilitate a directional membrane-mediated interaction between I-BAR domains.

## Introduction

Membrane curvature is generated throughout the cell and is required for many cellular functions. The Bin/Amphiphysin/Rvs (BAR) domain superfamily regulates membrane shape in a range of cellular processes ranging from endocytosis to filopodia formation.(1–4) Across the superfamily, full-length BAR proteins may have actin-binding domains, specific lipid targeting domains, or terminal amphipathic helices that are key to the proper function and localization of BAR proteins.(5) Notably, BAR domains share a consistent dimeric structure of bundled, kinked helices, and net positive charge. Among the many factors that distinguish BAR domain function are the membrane-binding interface and protein intrinsic curvature. The three families comprising the BAR superfamily are the N-, F- and I-BAR families: N- and F-BAR domains have similar concave membrane-binding interfaces while I-BAR domains are distinct in their convex binding surface. Subsequently, the function of I-BAR proteins is dissimilar as it is responsible for forming extracellular membrane protrusions (e.g., filopodia) instead of forming cytosolic membrane structures.(1)

Experimentally, the mechanism by which I-BAR domains induce membrane curvature has been interrogated by *in vivo* and *in vitro* studies. *In vivo* experiments show that I-BAR domains are essential to protrusion formation and the over expression of I-BAR proteins results in the formations of membrane protrusions from the cell.(6) *In vitro* experiments show I-BAR domains will sort into membrane tubules of a preferred curvature, as well as cause membrane deformation and tubulate giant unilamellar vesicles (GUVs).(7, 8) The onset of tubulation has been shown to be affected by membrane composition with phosphatidylinositol 4,5-bisphosphate (PIP_2_) playing an important role in lowering the necessary surface density of I-BAR domains required to induce GUV shape change.(9) Indeed, mutagenic and fluorescence studies have shown that I-BAR domains can cluster negatively-charged PIP_2_ due to the positively-charged residues present on the ends of I-BAR domains.(6, 10, 11)

Various computational and theoretical methods agree that a concerted deformation of many I-BAR domains gives rise to large-scale shape change. At an atomistic resolution, multiple I-BAR domains are difficult to simulate due to the computational burden of modeling the entire I-BAR domain, lipid membrane, and solvent environment. As such, only isolated I-BAR domains have been studied at atomistic resolution and, importantly, these simulations show that I-BAR domains have smaller intrinsic curvature when bound to the membrane as compared to predictions from their crystal structure.(12–15) Separately, particle-based coarse-grained (CG) models have been used to understand the membrane-bound behavior of many I-BAR domains.(15, 16) Notably, the aggregation and collective membrane deformation behavior of I-BAR domains is directly dependent on the local curvature induced by a single protein.(17, 18) Simpler field theoretic models have shown that I-BAR domain organization inside of membrane tubules should be expected due to geometric constraints of curved proteins in tubules.(19) Additional studies related to coupling of membrane and protein curvature elucidate the mechanism of membrane-mediated attraction that depends on membrane mechanical properties.(20, 21) While these studies differ in protein and membrane representations, they have elucidated how the membrane can give rise to membrane-mediated forces.(22–24) However, the role of membrane composition, and in particular, negatively-charged lipids such as PIP_2_, has not been investigated at this scale, such that the role of lipid clustering on protein-protein attraction remains unclear.

In this study, we aim to better understand membrane-mediated forces involved in I-BAR domain aggregation and membrane remodeling. We introduce highly CG membrane and protein models. We characterize the phenomenological aggregation behavior and membrane protrusion stabilization of I-BAR domains by investigating the effects of I-BAR domain curvature, protein-membrane interaction strength, and membrane composition. Taken together, these results help us to gain further insight into the membrane-mediated forces that drive I-BAR protein aggregation and stabilization of membrane protrusions.

## Methods

The membrane model is a solvent-free CG model recently developed by Grime and Madsen.(25) The model represents the membrane patches by three beads with tunable fluidity, rigidity, and intrinsic curvature as modulated by the inter- and intra-membrane patch interactions. The membrane-membrane and membrane-protein interactions follow the form shown in Equation 1, which is a soft, sine-based pair potential.

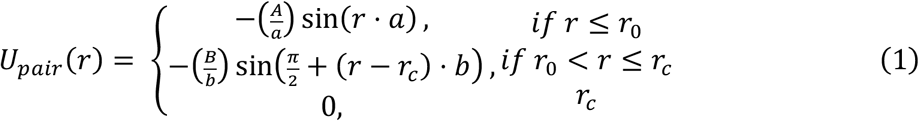

The membrane-membrane interactions of the upper and lower beads (purple CG beads in Figure 1A) are purely repulsive (i.e., *B* value of 0 *k*_B_*T*) and, in this study, both upper and lower CG beads have an *r_0_* value of 1.125 nm, resulting in no intrinsic curvature. The center bead (grey beads in Figure 1A) has an interaction strength or *B* value of 9.0 *k*_B_*T* to other center beads and *B* value of 0 *k*_B_*T* to other membrane beads and a *r_0_* value of 1.5 nm to all beads. The membrane parameter set results in a fluid monolayer with a bending modulus of ~10 *k*_B_*T*. The bonded interactions are harmonic with zero energy distance of 1.5 nm and spring constant of 25 *k*_B_*T*/Å^2^ or ~15 kcal/mol/Å^2^. The angular potential is harmonic with zero energy angle of 180 degrees and spring constant of 10 *k*_B_*T*/Å^2^ or ~6 kcal/mol/Å^2^. At this CG resolution, the membrane beads do not correspond to individual lipids, but effectively represent a patch of the lipid bilayer. From the perspective of simpler field theoretic models, the membrane acts as an undulating sheet that mediates protein-protein interactions via curvature coupling and Casimir-like forces. The highly CG nature of the membrane allows for large system sizes necessary for the simulation of the collective behavior of proteins at near micron length scales.

**Figure 1:**
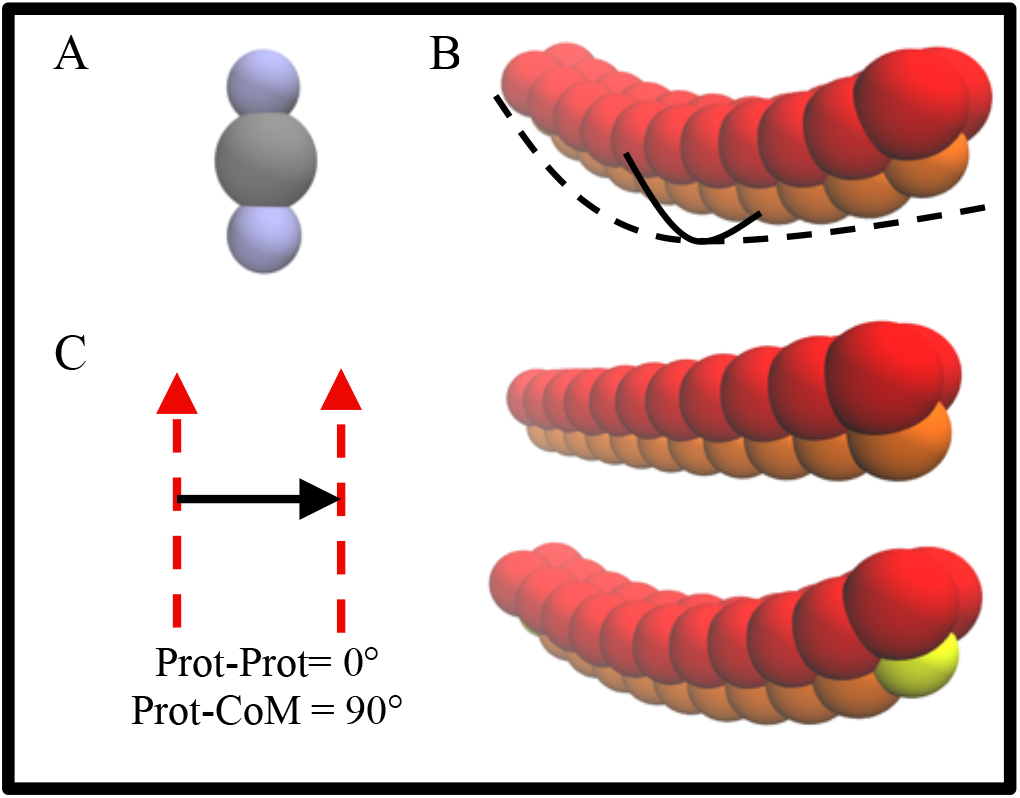
Membrane presentation (A) and three protein representations with first, curved I-BAR domain with uniform binding surface (orange) (B upper), flattened I-BAR domain with uniform binding surface (B center), and curved I-BAR domain with negatively-charged lipid binding site (yellow) (B lower) highlighting the transverse (solid) and longitudinal (dashed) curvatures. Schematic of two angles used to characterize protein aggregates: the angle formed by vectors describing the long dimension of two neighboring proteins (Prot-Prot) and the angle formed by a vector describing the long dimension of a protein and the center of mass between two neighboring proteins (Prot-CoM).

The protein model is represented by three linear strings with tunable longitudinal and transverse curvatures. The intra-protein interactions are an elastic network holding the protein rigid. The width and length of each I-BAR domain is consistent with the dimensions from the crystal structure of the I-BAR of IRSp53, while the curvature ranges from 1/15 nm^−1^ to 1/50 nm^−1^ (i.e., nearly flat).(12, 13) The protein-protein interactions are of the same form as the membrane-membrane interactions shown in Equation 1 and are purely repulsive (i.e., B value of 0) and *r_0_* of 1.7 nm.

The protein-membrane interactions are purely repulsive except in the case of the center string of I-BAR domain beads (see orange beads in Fig. 1B) and the middle monolayer bead (see grey beads in Fig. 1A). These interactions are varied from having an interaction strength of ~0.6 to ~3.2 kcal/mol, which effectively modulates the binding energy of the I-BAR domain and the local curvature induced by an individual I-BAR domain. By spanning a range of membrane-protein attractions, we assess the onset of protein clustering and characterize the resultant I-BAR domain clusters, which, for example, implicitly include membrane-protein electrostatic interactions. Additionally, I-BAR domains commonly have several positively-charged residues at the ends of the curved structure.(11) Thus in select cases, the end CG beads of the I-BAR domain (see yellow beads in Fig. 1B) have specific attraction to a subset of the membrane CG beads, e.g., to represent selective binding to PIP_2_-containing lipid patches. We modulate the membrane binding surface of the protein to probe how attractive membrane bead clusters affect aggregation behavior of I-BAR domains and the protrusion stabilization.

We ran these simulations using the LAMMPS molecular dynamics simulator (26) with a timestep of 200 fs using the Langevin thermostat (27) with temperature dampening of 20 ps and an anisotropic Parrinello-Rahman barostat (28) with pressure dampening of 200 ps. The periodic flat sheet simulations fluctuate around 160 nm by 160 nm with 44,409 membrane beads. The larger protrusion simulations contained 188,031 CG membrane beads with an initial lateral dimension of 300 nm that varies based on the radius of the protrusion made. The protrusions were formed using an external time-dependent repulsive cylindrical and spherical potentials implemented in LAMMPS to enforce a minimum protrusion radius. The walls were described using a 9-3 Lennard-Jones form with an epsilon value of 0.1 kcal/mol and sigma value of 30.0Å and deformed the membrane to form a 130nm protrusion over the course of a 10 μs simulation. The I-BAR domains coated the protrusion and were equilibrated by ramping the protein-protein repulsion down to 5 or 10% and back up, which facilitates equilibration of high-density coats of proteins. We ramped the repulsion down, ran with weak repulsion, and ramped back three times over the course of 16.2 μs. Then, we ran with full repulsive interactions for 5 μs, which were analyzed for I-BAR domain statistics in protrusions formed by external potentials. During this time, the protein aggregate can deform the shape of the tubule outward, away from a cylindrical protrusion, because the external potential only enforces a minimum protrusion radius. Finally, the external bias is removed, and both the membrane and proteins are relaxed for another 25 μs. We use the last 5 μs for statistics of relaxed protrusions.

We quantified I-BAR domain aggregation using two characteristic angles shown in Figure 1C: The angle between two adjacent proteins (Prot-Prot) and the angle formed by the protein and the center-of-mass vector between two adjacent proteins (Prot-CoM). We characterized the stability of protrusions by calculating the local curvature over the entire protrusion. We calculated local curvature by clustering membrane beads, performing a local dimensionality reduction with the isomap algorithm, next performing a Delaunay triangulation in the reduced dimensions and, finally, using the triangles in the full three dimensions to calculate the curvature at each cluster center.(29) All characterization was done in Python using numpy,(30) scipy,(31–33) and scikit-learn(34).

## Results

### Flat sheet organization as a function of binding strength and protein curvature

Initially, we ran flat sheet simulations of the I-BAR domains to explore and characterize the aggregation behavior of the current I-BAR domain model in comparison to previous particle-based BAR domain models. We found that I-BAR domain curvature and binding strength, as modulated by the membrane-protein interaction strength, control the aggregation behavior. At I-BAR domain radii greater than 30 nm (curvature less than 0.033 nm^−1^) and interaction strengths below 1.9 kcal/mol, the proteins do not aggregate (see black lines in Fig. 2A). As membrane-protein interaction strength increases, the I-BAR domains form end-to-end linear aggregates (see yellow squares in Fig. 2A).(24) At low interaction strength and increasing curvature, the I-BAR domains form partial side-by-side aggregates. At high curvature and increasing interaction strength, the preferred aggregate morphology changes from end-to-end, through an intermediate aggregate of side-by-side/end-to-end aggregate and, eventually, side-by-side aggregates (see purple circles in Fig. 2A). By tuning the protein-membrane interactions and protein curvature, we show a rich phenomenology of aggregation behavior similar to previous models of I-BAR domains.(15, 16) This provides further evidence that membrane-mediated forces (i.e., curvature-mediated and Casimir-like attraction) are sufficient to drive membrane aggregation in the absence of direct protein-protein attraction. Next, we introduced an additional membrane-mediated force due to attractive membrane bead clustering.

**Figure 2:**
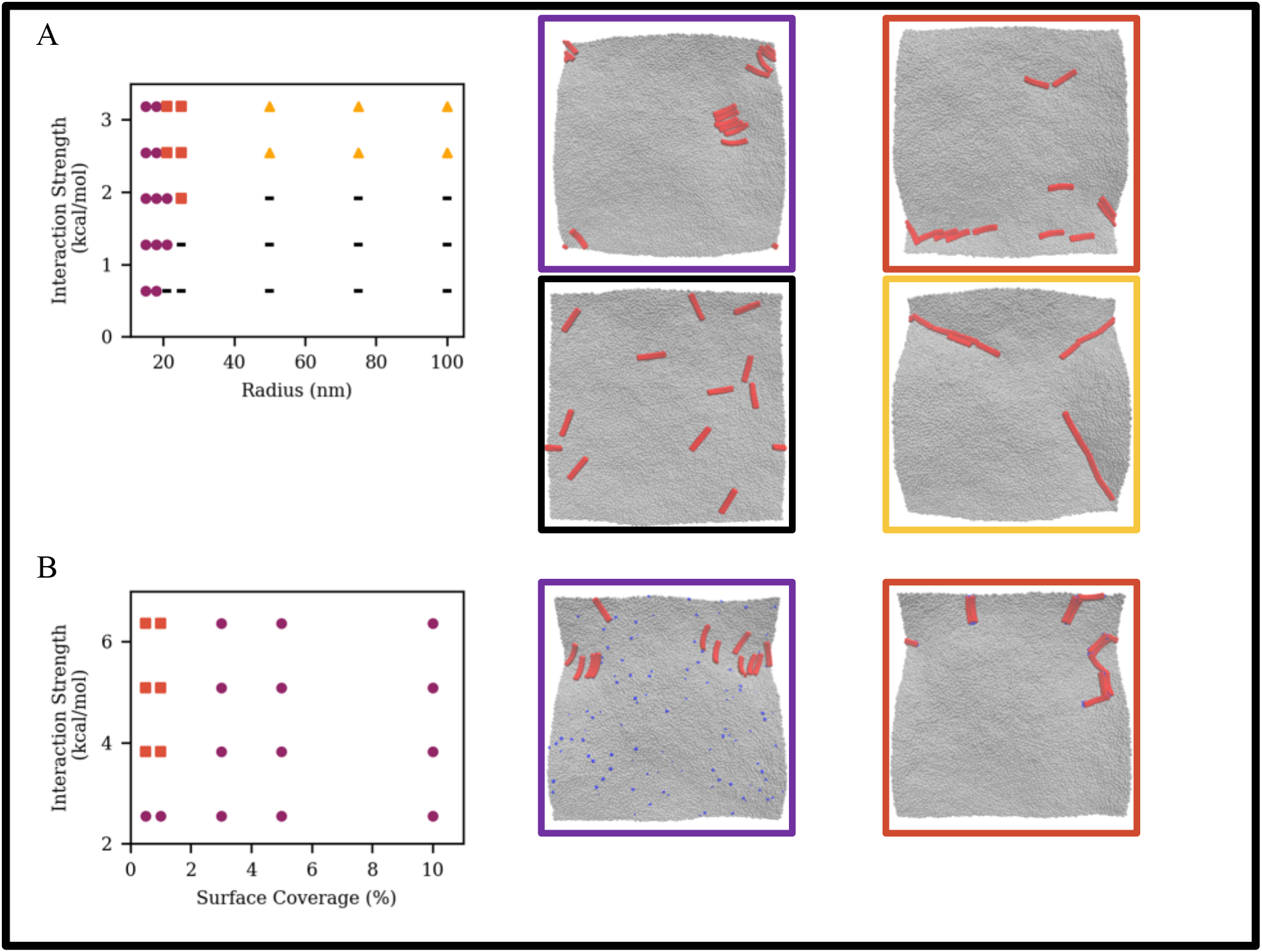
Characterization of aggregation behavior as a function of protein curvature and protein-membrane interaction strength without PIP_2_-like membrane beads (A) and as a function of PIP_2_-like membrane interaction strength to I-BAR binding site and PIP_2_-like surface coverage for 18nm radius I-BAR domain (B). Representative snapshots of no aggregation (black lines), end-to-end aggregate (yellow triangles), intermediate aggregates (red squares), and side-by-side aggregate (purple circles),

### Flat sheet organization with attractive membrane beads

We sought to understand the role of lipid clustering and how it might give rise to a new local membrane-mediated force. To capture this effect, e.g., for PIP_2_ clustering, we made two changes to our low-resolution CG model. We introduced a PIP_2_-like membrane bead that has stronger attraction to the ends of I-BAR domains shown in yellow in the lower panel of Fig. 1B, thereby making the membrane binding surface of the I-BAR domain non-uniform. The non-uniform membrane binding surface is physically motivated by the higher density of positive charge near the ends of I-BAR domains due to a large number of positively-charged residues located at the ends. Thus, the non-uniform CG interaction surface can recapitulate the preferred interactions between the ends of an I-BAR domains and a subset of the membrane beads. Additionally, we chose a protein curvature of 1/18 nm^−1^ and protein-membrane interaction strength of 0.6 kcal/mol for the rest of the membrane-binding surface. Then, we characterized the aggregation behavior as a function of the I-BAR domain-attractive membrane bead interaction strength and the effective surface coverage of attractive membrane beads.

As expected, we found that attractive membrane beads cluster to the ends of the I-BAR domains. We found that the highly attractive membrane patch significantly affects the I-BAR domain organization by introducing an area where I-BAR domains can co-localize. In the case of curved I-BAR domains, the side-by-side aggregates can be destabilized, and end-to-end aggregates preferred because the I-BAR domains organize around a central attractive membrane bead (see red squares of Figure 2B).

In Figure 2, we show snapshots of the 18 nm radius (or 0.056 nm^−1^ curvature) I-BAR domain with two sites to bind attractive lipids while the rest of the membrane binding surface is unchanged from previous simulations. These choices are used to closely model the membrane binding surface of I-BAR domains that have several positively-charged residues at the ends. The flat sheet aggregation behavior changes drastically with the addition of attractive membrane beads. The attraction strength between the ends of I-BAR domains to the PIP_2_-like lipid patches was taken as a free parameter, which we investigated from 2.5 kcal/mol to 6.4 kcal/mol. When the interaction strength between the end of the I-BAR domain and the attractive membrane beads is five or six times stronger than other membrane beads (i.e., interaction strength greater than 10), there is an onset of clustering and the aggregation behavior is affected (see red squares in Fig. 2B). When there is a uniform binding surface, we found the 18nm radius I-BAR domain with weak membrane-protein interactions would form weak side-by-side aggregates. As we added PIP_2_-like membrane beads (i.e., the surface coverage of the PIP_2_-like lipids increases), we found that both the side-by-side and end-to-end aggregates are stabilized. Furthermore, as we increased the surface coverage of attractive membrane beads, we find that end-to-end aggregates are less probable than side-by-side aggregates.

### Protrusion stability

Next, we probed the stability of the membrane protrusions coated with I-BAR domains. We formed a protrusion using an external potential, removed the external potential, and allowed the protrusion to relax under the influence of only I-BAR domain coats. The diameter of the protrusion was small compared to the lateral dimension of the simulation box (e.g., 30 nm diameter protrusion is formed in a 300 nm box). We swept over a range of protein and protrusion curvatures; Figure 3 shows representative snapshots for 15, 21, and 50 nm radius I-BAR domains (left, center, right columns of Figure 3A and B) inside of 15 and 21 nm radius protrusions as well as the characterization of membrane Gaussian curvature and protein number density. The snapshots omit the large flat region surrounding the protrusion. Over this range of radii, we found a variety of relaxed protrusion configurations. When comparing the left and center column of Figure 3A, we saw that while the protrusion radius is initially 15nm, after relaxation the center protrusion is wider corresponding to the radii of the proteins inside. Generally, we find that the curved I-BAR domains change the radius of a protrusions towards the radius of the I-BAR domains, and flat I-BAR domains did not stabilize the protrusions and sort toward the flat area surrounding the protrusions.

**Figure 3:**
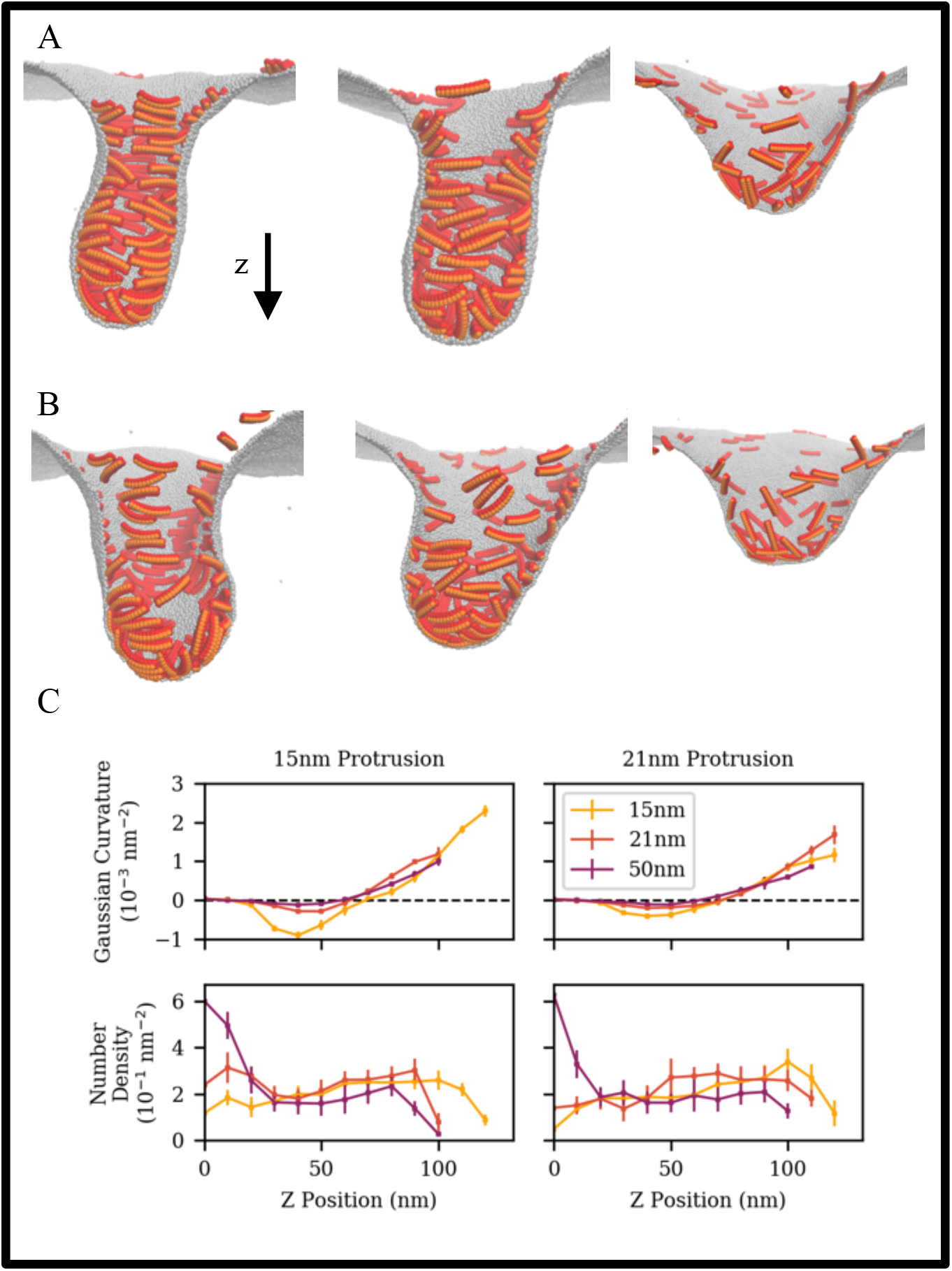
Snapshots of protrusions of 15nm (A) and 21nm (B) radii stabilized by I-BAR domains of 15nm, 21nm, and 50nm (left to right) radii. Gaussian curvature and number density as position along the length of the protrusion (C) for the corresponding protrusion and I-BAR domain radii.

In the case of 15 nm I-BAR domains inside of 15 nm radius protrusions, aggregates stabilize a bulbous protrusion (see Fig. 3A left). Indeed, when we calculate the Gaussian curvature profile along the length of the protrusion, the protrusion does not have a region of 0 curvature, which is a characteristic of cylinders. We do see constantly varying Gaussian curvature along the length of the protrusion. Additionally, we see a relatively flat number density of I-BAR domains meaning there is little sorting in or out of the protrusion. Inside of a wider, 21 nm protrusion (see Fig. 3B left), the I-BAR domain aggregate stabilizes the bulbous protrusion with constantly varying Gaussian curvature of a lower magnitude as compared to the 15 nm protrusion. In the intermediate case of 21 nm I-BAR domains (see Fig. 3A center), we find that the stabilized protrusion is similarly bulbous as well as widened by the I-BAR domain aggregate towards the radius of the I-BAR domain. When the I-BAR domain radius and protrusion radius are both 21 nm, the stabilized protrusion is similar to the 15 nm protrusion case. Although, we do find that the number density on the flat sheet to be slightly lower when the initial protrusion is 21 nm as compared to 15 nm, corresponding to a weak sorting effect in the 21 nm protrusion. In the case of low curvature I-BAR domains (i.e., flat I-BAR domains) shown in right snapshots of Figure 3A and 3B, the protein aggregate is not sufficient to stabilize the protrusion. Indeed, when considering the Gaussian curvature and number density profiles, we find that there is little to no negative Gaussian curvature and the I-BAR domains have a significantly higher density on the flat membrane surrounding the protrusion.

We compared protrusion stabilization in the presence of attractive membrane patches at the I-BAR ends. We found that the I-BAR domain aggregates continue to cluster the attractive membrane beads in between side-by-side aggregates (See Fig 4A). When comparing Gaussian curvature profiles shown in Figure 4C, protrusions with PIP_2_-like membrane beads have a small but distinct shoulder in the Gaussian curvature profile, while protrusions without PIP_2_-like membrane beads do not. In other words, protrusions with the attractive membrane beads are more cylindrical than those without. The attractive membrane beads mediate an attraction between side-by-side aggregates, constricting the protrusion towards the curvature of the I-BAR domains. Thus, we show that PIP_2_-like membrane beads serve a key functional role in stabilizing cylindrical membrane protrusions, and this is one of the primary conclusions from the present work.

**Figure 4:**
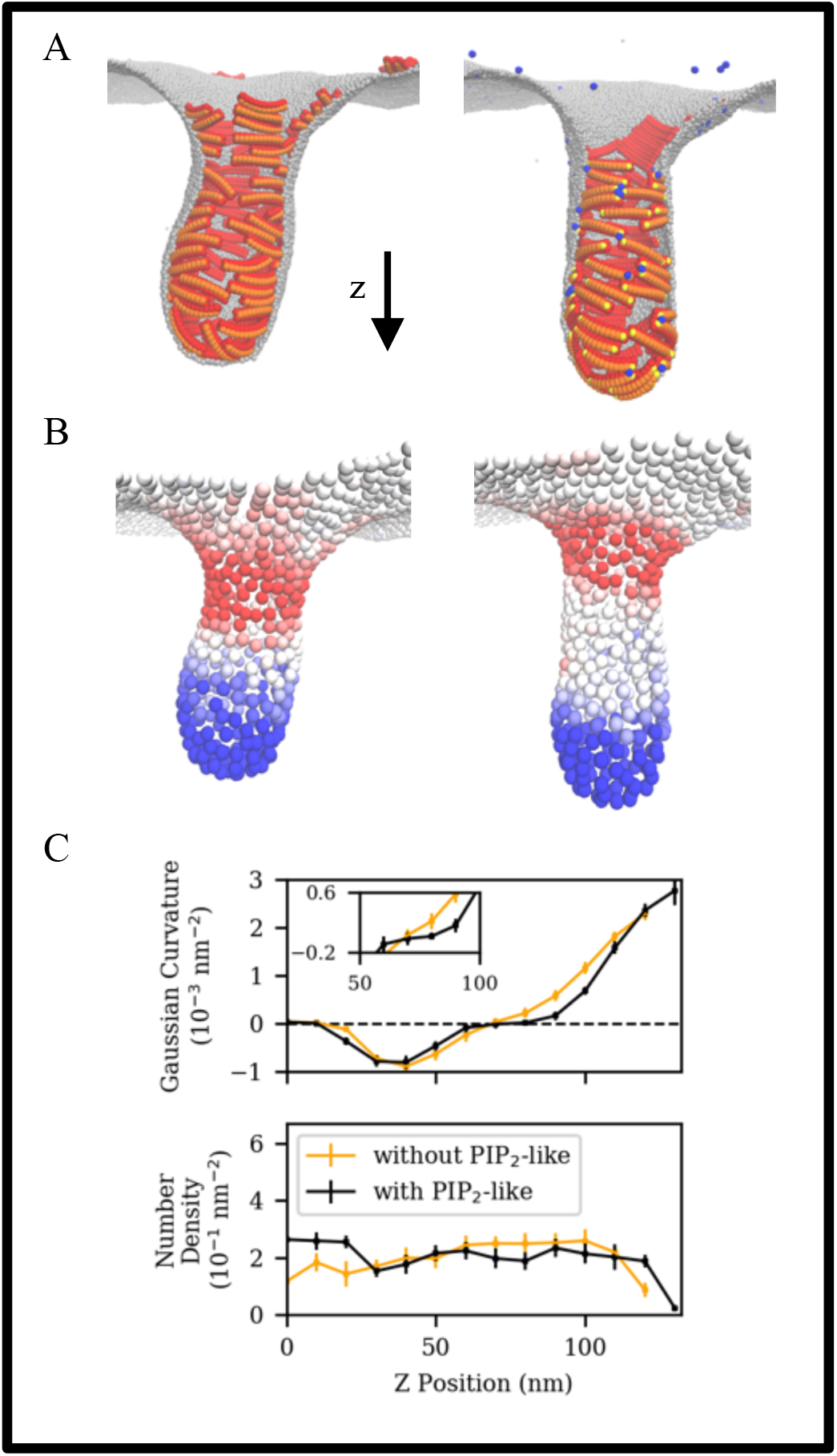
Snapshots of protrusions stabilized by 15nm I-BAR domains with and without PIP_2_-like membrane bead (A: left and right, respectively), the reduced representation of membrane (B: negative and positive Gaussian curvature shown in red and blue, respectively), and Gaussian curvature and number density as position along the length of the protrusion (C).

## Discussion

### Role of PIP_2_-like membrane regions in flat sheet organization

Our goal was to determine the essential physics of I-BAR domain-mediated membrane remodeling. The generic nature of our model captured only the key characteristics of membrane and the I-BAR domain family. We began by accounting for shape, binding energy of I-BAR domains, and the deformation energy of the membrane. We showed that our model can express a variety of organization and aggregation behavior that is consistent with previous BAR domain models using strictly pair-wise interactions. As the interactions between I-BAR domains are completely repulsive, membrane-mediated attraction is attributed to curvature-inducing and thermal fluctuation Casimir-like forces. Indeed, when membrane-protein interactions are weak, the I-BAR domain induces little to no local membrane deformation and does not aggregate. When protein curvatures and membrane-protein interaction strengths are varied, we observe a variety of side-by-side and end-to-end I-BAR domain aggregates. The end-to-end aggregates are curvature-driven primarily due to the transverse curvature of the I-BAR domain. Each flattened protein generates local deformations in the form of a membrane trough and the proteins align to create a single long membrane trough similar to the behavior seen in previous bottom-up CG model of I-BAR domain.(15) As the curvature along the long-dimension of the protein increases, the curvature coupling in the long dimension dominates and end-to-end aggregates give way to more stable side-by-side aggregates. When side-by-side aggregates form, we find that there is an increase in membrane curvature, but in our flat sheet simulations, we do not find any tubulation.

The absence of tubulation suggests that our model does not include CERTAIN details that are required for membrane tubulation. For example, there may be direct protein-protein interactions, or a more complex membrane binding surface (e.g., specific PIP_2_ binding) not being taken into account in our generic model. We hypothesize the competition between side-by-side and end-to-end aggregates to be important for the onset of tubulation, as we found solely side-by-side or end-to-end aggregates to be insufficient to induce tubulation.

### Role of PIP_2_-like, attractive membrane regions in tubule stabilization

We introduced an attractive membrane CG bead meant to capture the effect of a PIP_2_-like lipid that is specifically attracted to the ends of the I-BAR domain, thereby representing an additional membrane-mediated force. After characterization of I-BAR domain aggregates, we find that the PIP_2_-like membrane beads significantly change the aggregation behavior on flat sheets. When comparing representative snapshots in Figure 2B, we find clustered PIP_2_-like membrane beads are located at the ends of the I-BAR domain causing a competition between end-to-end and side-by-side aggregates at low surface coverages of PIP_2_-like membrane beads.

When modeling protrusion stability, we noted that the protein aggregation behavior inside of tubules is directly affected by the addition of PIP_2_-like, attractive membrane beads. When the protein has a uniform membrane binding surface, curved proteins will form side-by-side aggregates inside of the protrusions. However, gaps emerge between side-by-side aggregates that give rise to irregularly shaped membrane protrusions with constantly varying Gaussian curvature. When we introduce the attractive membrane beads and relatedly, the membrane binding surface, the PIP_2_-like membrane beads cluster and serve to co-localize side-by-side aggregates, which collectively induce membrane tubule constriction. Ultimately, the presence of PIP_2_ appears to yield protrusions with more regular cylindrical shapes and reduced membrane curvatures. We thus find that PIP_2_-like membrane beads provide a new membrane-mediated driving force that helps stabilize cylindrical protrusions. We hypothesize that the membrane-mediated interaction gives rise to the experimental observation that PIP_2_ reduces the amount of bound I-BAR domains necessary to induce significant shape change.(9)

### Connection to experiments

We directly investigated the sensitivity of protein aggregation and tubule stabilization across a wide range of physically-relevant parameters. We consider this an advantage of phenomenological CG model. We now aim to connect various parameters of the model to experimentally measurable quantities. For example, the membrane parameters were adjusted such that the membrane bending rigidity is consistent with that of *in vitro* membranes.(7) Another key parameter was the protein curvature, which we varied to reflect the diversity throughout the I-BAR domain family and the flexibility seen in atomistic simulations.(11, 14, 15) As we investigated protein curvature, we found that proteins with insufficiently small curvature fail to stabilize protrusions, which would be an interesting hypothesis to test experimentally. Finally, we note that the external potentials used to initiate tubule formation are conceptually similar to the force applied in tube pulling assays. In these experiments, the membrane tubules are not generated spontaneously by I-BAR domains, but are formed by beads pulled by pipettes or optical traps, allowing the I-BAR domains to sort into the tubules.(35, 36) Analogously, the protrusions in our simulations are initially generated by an external potential and the I-BAR domains sort into/out of the tubules depending the protein and protrusion curvature. Our results show protein sorting as flattened proteins have higher densities outside of the protrusion than curved proteins.

The strength of membrane-protein interactions is another key parameter that our simulations identified as a modulator for protein aggregation behavior. We probe the nonspecific attraction to the membrane as well as the specific attraction to a PIP_2_-like membrane bead. Experimentally, both specific and nonspecific interactions may be tuned by adjusting electrostatic forces between I-BAR domains and the membrane. The specific interactions could be tuned by changing the location of positively-charged residues in the I-BAR domain sequence to change the charge density of the membrane binding surface, while the nonspecific interactions could be tuned by varying the membrane composition with fewer anionic lipids, varying salt content, or mutating the I-BAR domain to change the total number of positively-charged residues. Thus, our findings provide intuition for future experiments investigating the role of lipid clustering and the membrane binding surface.

## Conclusions

Our phenomenological model shows the stabilization of membrane protrusions by an I-BAR domain interaction mediated by clustering lipids to the ends of I-BAR domains. We demonstrated I-BAR domain aggregation and organization of flat membranes is significantly different when PIP_2_-like membrane patches are present. When I-BAR domains have a uniform membrane binding surface, curved I-BAR domains form side-by-side aggregates. When I-BAR domains have preferred lipid binding domains, curved I-BAR domains form a variety of aggregates due to competition between membrane-mediated forces (i.e., forces due to curvature induction, thermal fluctuations and lipid clustering). Thus, we show that membrane composition directly affects membrane aggregation resulting in differing membrane protrusion stabilization.

### Author Contributions

Z.J and G.A.V. designed research; Z.J. performed research; Z.J., P.B. and G.A.V. analyzed data; Z.J., P.B. and G.A.V. wrote the paper.

## Acknowledgements

This research was supported in part by the National Institute of General Medical Sciences of the United States National Institutes of Health under NIH award number R01-GM063796 (Z.J and G.A.V.), and in part by the Human Frontiers Science Program through grant RGP0005/2016 (P.B., Z.J., and G.A.V.)

